# Long-term meditation training is associated with enhanced attention and stronger posterior cingulate – rostrolateral prefrontal cortex resting connectivity

**DOI:** 10.1101/2021.09.14.460351

**Authors:** Tammi RA Kral, Regina Lapate, Ted Imhoff-Smith, Elena Patsenko, Daniel W Grupe, Robin Goldman, Melissa A Rosenkranz, Richard J Davidson

**Affiliations:** Center for Healthy Minds, University of Wisconsin – Madison; Waisman Laboratory for Brain Imaging and Behavior, University of Wisconsin – Madison; Department of Psychology, University of Wisconsin – Madison; Department of Psychiatry, University of Wisconsin – Madison; Department of Psychological and Brain Sciences, University of California, Santa Barbara

## Abstract

Mindfulness meditation has been shown to increase resting state functional connectivity (rsFC) between the posterior cingulate cortex (PCC) and dorsolateral prefrontal cortex (DLPFC), which is thought to reflect improvements in attention to the present moment. However, prior research in long-term meditation practitioners lacked quantitative measures of attention that would provide a more direct behavioral correlate and interpretational anchor for PCC–DLPFC connectivity and was inherently limited by small sample sizes. Moreover, whether mindfulness meditation primarily impacts brain function locally, or impacts the dynamics of large-scale brain networks, remained unclear. Here, we sought to replicate and extend prior findings of increased PCC – DLPFC rsFC in a sample of 40 long-term meditators (average practice= 3759 hours) who also completed a behavioral assay of attention. In addition, we tested a network-based framework of changes in inter-regional connectivity by examining network-level connectivity. We found that meditators had stronger PCC-rostrolateral prefrontal cortex (PFC) rsFC, lower connector hub strength across the default mode network (DMN) relative to other functional networks, and better attention to task, compared to 124 meditation-naïve controls. Orienting attention positively correlated with PCC– rostrolateral PFC connectivity, and negatively correlated with DMN connector hub strength. These findings provide novel evidence that posterior cingulate – rostrolateral PFC rsFC may support attention orienting, consistent with a role for rostrolateral PFC in meta-cognitive background awareness that is a core component of mindfulness meditation training. Our results further demonstrate that long-term mindfulness meditation may improve attention and strengthen the underlying brain networks.

## Introduction

Mindfulness meditation is defined as the practice of focusing attention on present-moment experience (Kabat-Zinn, 1990), in contrast to mind-wandering or attending to thoughts about the past or future. A growing body of research suggests that mindfulness practice is associated with alterations in activation and connectivity of the posterior cingulate cortex (PCC) (Brewer and Garrison, 2014), a region that is part of the ‘default mode network’ and that has been previously implicated in mind wandering (Fox et al., 2015; Spreng et al., 2008). Indeed, participants had increased PCC resting state functional connectivity (rsFC) with dorsolateral prefrontal cortex (DLPFC) following a short-term Mindfulness-Based Stress Reduction (MBSR) intervention compared to an active control group (Kral et al., 2019). Moreover, the same study found that increased PCC –DLPFC connectivity was associated with improved attention. However, prior research on differences in PCC rsFC among long-term mindfulness meditation practitioners lacked quantitative measures to clarify the behavioral relevance of PCC – DLPFC connectivity (Brewer et al., 2011).

A recent meta-analysis of functional neuroimaging studies of meditation practice found that focused attention meditation – an essential ingredient of mindfulness practice – reduced PCC activation (Fox et al., 2016). De-activation of PCC was associated with “undistracted awareness” and “concentration” based on a qualitative analysis of meditators’ subjective reports during an fMRI neurofeedback task (Garrison et al., 2013). Moreover, long-term meditators were able to purposefully deactivate PCC through meditation during neurofeedback, whereas meditation-naïve control participants were unable to do so (Garrison et al., 2013). Therefore, PCC likely plays a central role in meditation-related improvements in attention and studies on long-term mindfulness practitioners may provide unique insights into the effects of mindfulness meditation on brain function and connectivity, shedding light on unique mechanisms of change as a function of specific stages of training (Brefczynski-Lewis et al., 2007; Kral et al., 2018).

Short- and long-term mindfulness meditation training have both been associated with increased rsFC between PCC and DLPFC (Brewer et al., 2011; Kral et al., 2019). The DLPFC is a key node of the frontoparietal control network associated with attentional control (MacDonald et al., 2000; Smallwood et al., 2012). Increased PCC – DLPFC coupling is thought to reflect better attentional control over mind-wandering (Brewer and Garrison, 2014). Indeed, prior research that employed a sub-sample of participants examined in the current study provided initial evidence in support of this hypothesis: changes in PCC – DLPFC connectivity were positively associated with increased self-reported attentional control following an 8-week MBSR program (Kral et al., 2019). However, prior research with long-term mindfulness meditation practitioners lacked measures of attention to test the hypothesis that increased PCC – DLPFC connectivity reflects improved attention, and was further limited by small sample sizes of less than fifteen participants per group (Brewer et al., 2011). Therefore, the functional relevance of increased PCC – DLPFC connectivity among long-term mediation practitioners remains unclear.

The current study builds on this literature by examining relationships between long-term mindfulness meditation practice and PCC rsFC in a sample of 40 meditators (vs. 124 meditation-naïve controls) who were later assigned to an intervention as part of a randomized controlled trial (RCT) (as detailed in Kral *et al*., 2019). Following prior work, we examined changes in PCC rsFC with 3 regions of interest (ROIs): left and right DLPFC, and left rostrolateral PFC. The DLPFC ROIs were anatomically defined based on the middle frontal gyrus (depicted in teal in Figure 1c), and overlapped with ROIs where long-term meditation practice was previously shown to relate to stronger rsFC with PCC (Brewer et al., 2011). Given the size and potential for functional heterogeneity within this DLPFC ROI, we conducted analysis of mean ROI connectivity, as well as voxelwise analysis within the DLPFC mask. The rostrolateral PFC ROI (depicted in green Figure 1c) was defined based on coordinates from a study showing a significant effect of mindfulness meditation training on PCC rsFC (Creswell et al., 2016), and was located rostral to the canonical DLPFC. We previously examined changes in PCC rsFC with this rostrolateral PFC ROI following MBSR in a sub-sample of participants from the current study (Kral et al., 2019), and the current study extends this analysis to test for effects of long-term mindfulness training. In addition, we tested for relationships between PCC rsFC and total lifetime home and retreat practice hours, separately, and hypothesized that more hours of lifetime meditation practice would be associated with a larger increase in PCC rsFC.

**Figure 1.**
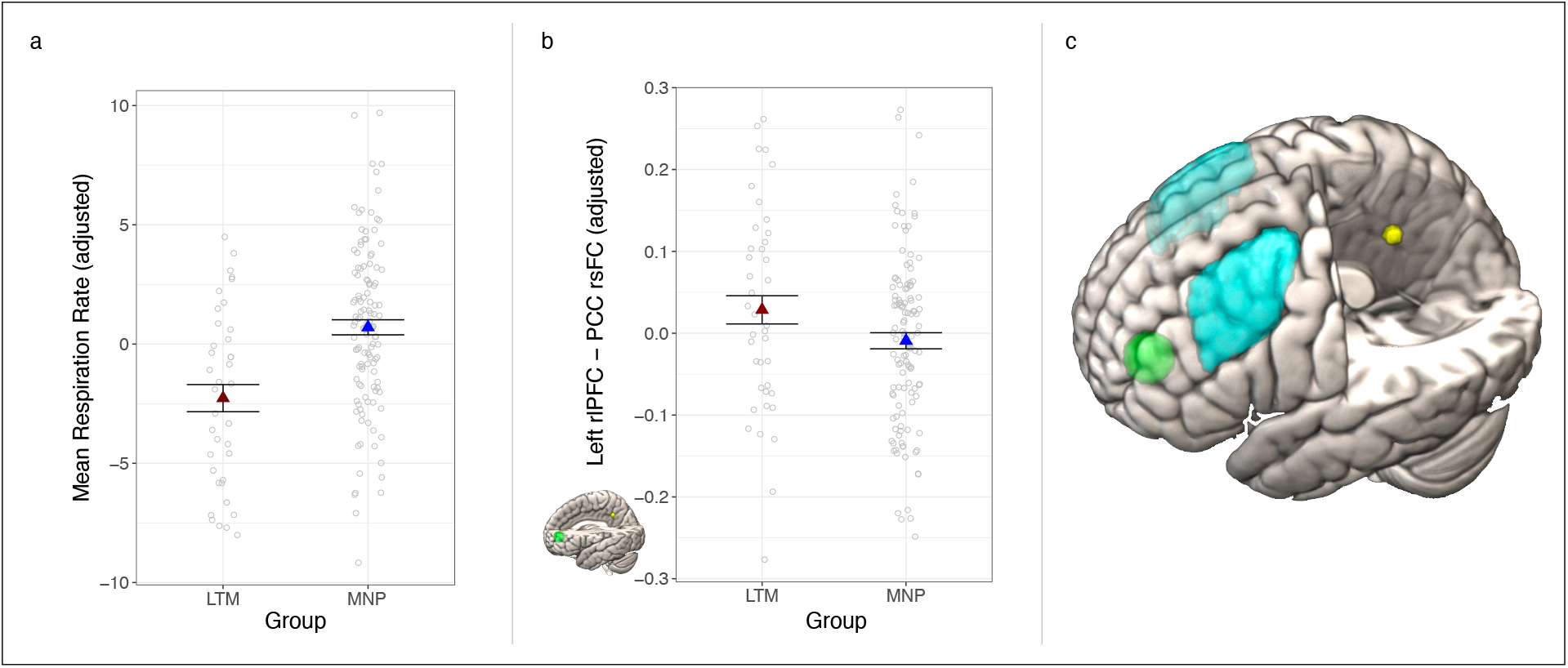
Long-term mindfulness meditation practice associated with slower respiration rate and stronger PCC-rlPFC rsFC. **(a)** Meditators had slower average respiration rate than non-meditators. **(b)** Meditators had stronger PCC rsFC with rostrolateral PFC (rlPFC) compared to meditation-naïve participants. The rlPFC target region is depicted in green, and the PCC seed region is depicted in yellow. **(c)** The dorsolateral prefrontal cortex (PFC) ROIs are in light blue (based on the middle frontal gyrus from the Harvard-Oxford atlas (Craddock et al., 2012)), the rostrolateral PFC ROI is depicted in green, and the PCC seed is in yellow (with the latter two based on coordinates from a prior study (Creswell et al., 2016). Dependent variables and data points are adjusted for age and gender, and an additional covariate for scan acquisition version in (b). Error bars represent 1 standard error above and below the point estimates of the means. PCC = posterior cingulate cortex; rsFC = resting state functional connectivity; LTM = long-term meditator; MNP = meditation naïve participant; rlPFC = rostrolateral prefrontal cortex

We expected to replicate prior results of increased PCC resting state connectivity with dorsolateral and rostrolateral PFC in this larger sample of 40 long-term meditators relative to a meditation-naïve, age-matched control group, which would comprise a substantial advancement over prior reports with limited sample sizes (e.g., less than 15 participants per group; Brewer et al., 2011). Since individual differences in physiology may affect fMRI measures (de la Cruz et al., 2019), and long-term meditation has been associated with lower respiratory rate (Wielgosz et al., 2016), we also tested for group differences in heart rate and respiration rate, and conducted sensitivity analysis to test whether group differences in rsFC persisted after controlling for these physiological measures. We also sought to extend prior research findings in two critical ways. First, informed by our prior work, we examined relationships between PCC connectivity and measures of attention, which is critical for interpreting rsFC differences between long-term meditators and controls. To that end, we included two self-report measures and one behavioral measure: the attention scale of the Emotional Styles Questionnaire (ESQ) (Kesebir et al., 2019) and experience sampling from text messaging, and the Attention Network Task (ANT). The ANT provided a well-validated, behavioral measure of three types of attention: alerting, orienting and executive control (Fan et al., 2009).

Second, we examined group differences in graph theoretical metrics indexing network topography. We assessed whether rsFC differences associated with mindfulness meditation are regionally specific to the PCC, DLPFC, and rostrolateral PFC seeds, or whether they may instead reflect underlying differences in the overall dynamics of separate networks to which these regions belong. To that end, we focused on two graph theoretical measures obtained from resting connectivity estimates: within-module degree and participation coefficient, which index within- and between-module hub properties, respectively. Within-module degree indicates high local connectivity of a given node to other nodes within the same module, while participation coefficient denotes the diversity of intermodular connections. More specifically, a provincial hub is a node with high within-module degree. In contrast, a connector hub has a high participation coefficient, and is posited to contribute to global intermodular integration (Rubinov and Sporns, 2010).

If higher PCC – rostrolateral PFC connectivity in meditators reflects a more general difference in network dynamics, such that these modules are more integrated, then we would expect to see higher participation coefficients for one or both of the corresponding networks (default mode network [DMN] and frontoparietal control network, respectively). Given the interpretation of higher PCC – rostrolateral PFC connectivity as reflecting increased attentional control by the frontoparietal control network on default mode network, we hypothesized that meditators would have higher participation coefficients than meditation-naïve participants in the frontoparietal control network, and lower within-module degree in the DMN. Additionally, we assessed hub properties for the dorsal attention network, in which DLPFC participates.

Thus, we here sought to (a) replicate prior research showing mindfulness meditation-related increases in PCC functional connectivity with dorsolateral and rostrolateral PFC, (b) extend the literature to determine whether such differences were associated with improvements in attentional measures, and (c) assess network connectivity metrics derived from graph theory.

## Results

Corrected *p*-values are indicated by *p*\*. We used a false discovery rate (FDR) correction to control for multiple comparisons for each family of tests (e.g., across 3 ROIs). Descriptive statistics for dependent variables by group are presented in Table 1.

**Table 1.** Descriptive statistics.

| Measure | Long-term meditators |  |  |  | Meditation-naïve participants |  |  |  |
| --- | --- | --- | --- | --- | --- | --- | --- | --- |
|  | *M* | *SD* | *Min* | *Max* | *M* | *SD* | *Min* | *Max* |
| R DLPFC-PCC rsFC (*r*) | 0.17 | 0.14 | -0.12 | 0.51 | 0.13 | 0.12 | -0.10 | 0.49 |
| L DLPFC-PCC rsFC (*r*) | 0.19 | 0.12 | -0.01 | 0.54 | 0.16 | 0.10 | -0.09 | 0.51 |
| L RLPFC-PCC rsFC (*r*) | 0.17 | 0.14 | -0.13 | 0.46 | 0.12 | 0.11 | -0.11 | 0.40 |
| Attention (ESQ) | 4.84 | 0.91 | 2.86 | 6.43 | 4.39 | 1.01 | 1.86 | 6.43 |
| Attention (ES) | 7.39 | 1.06 | 4.72 | 8.96 | 6.68 | 1.09 | 3.57 | 9.00 |
| ANT-R orienting RT (ms) | 93.2 | 42.0 | -0.27 | 192.8 | 101.0 | 40.1 | 9.21 | 206.7 |
| ANT-R executive RT (ms) | 135.5 | 41.3 | 38.5 | 270.0 | 137.4 | 41.6 | 8.8 | 307.4 |
| ANT-R alerting RT (ms) | 23.0 | 25.5 | -33.6 | 76.0 | 19.4 | 26.6 | -41.9 | 117.3 |

R = right; L = left; DLPFC = dorsolateral prefrontal cortex; PCC = posterior cingulate cortex; rsFC = resting state functional connectivity; RLPFC = rostrolateral prefrontal cortex; ESQ = Emotional Styles Questionnaire; ES = experience sampling; SLF = superior longitudinal fasciculus; DTI = diffusion tensor imaging; ANT-R = Attention Network Task-Revised; RT = reaction time; ms = milliseconds; M = mean; SD = standard deviation; Min = minimum; Max = maximum

### Physiological measures

Group differences in respiration rate and heart rate may contribute to or confound rsFC differences between meditators and meditation-naïve participants (Chen et al., 2020; de la Cruz et al., 2019) and previous research has reported reduced respiration rate in long-term meditators (Wielgosz et al., 2016). Therefore, we examined group differences in respiration and heart rates in this sample. Meditators had significantly slower respiration rate than meditation-naïve participants (*t*(157)=4.60, *p*<0.001, *b*=3.04, CI=[1.73, 4.34]; Figure 1a), consistent with prior research. There was no difference in heart rate between meditators and meditation-naïve participants (*t*(148)=-0.46, *p*=0.64, *b*=-0.70, CI=[−3.68, 2.28]).

### Resting State Brain Connectivity

#### ROI analysis

We tested for differences in PCC rsFC between meditators and meditation-naïve participants using a linear model in which we regressed average rsFC Fisher-Z transformed *r*-values (from DLPFC and rostrolateral PFC ROIs) on group, including covariates for age, sex, and scan resolution (which was increased partway through data collection to better address sinus-related artifacts. See Methods for additional details). There were no differences in PCC rsFC between meditators and meditation-naïve participants with the anatomically-generated DLPFC ROIs (right DLPFC: *t*(155)=-1.57, *p*=0.12, *p*\*=0.18, *b*=-0.04, CI=[−0.08, 0.01]; left DLPFC: *t*(157)=-0.79, *p*=0.43, *p*\* 0.43, *b*=-0.02, CI=[−0.05, 0.02]). Thus, we failed to replicate prior findings of increased PCC-DLPFC functional connectivity associated with long-term meditation practice (Brewer et al., 2011).

Meditators had stronger rsFC than meditation-naïve participants between PCC and left rostrolateral PFC (*t*(154)=-2.78, *p*=0.01, *p*\*=0.03, *b*=-0.06, CI=[−0.10, −0.02]; Figure 1b). These results remained significant when we added respiration rate and heart rate to the model, and are consistent with prior work showing increased PCC – rostrolateral PFC connectivity following a short-term mindfulness meditation intervention (Creswell et al., 2016), but which were not replicated in the RCT with control participants from the current study (Kral et al., 2019). Subsequent ROI analyses focused on the rostrolateral PFC ROI in which we found significant group differences.

#### Network analysis

We next examined network-level connectivity measures in order to test whether long-term mindfulness meditation training was associated with connectivity differences in the larger brain networks in which PCC, DLPFC, and rostrolateral PFC are embedded. We regressed participation coefficients and within-module degrees (separately) for all nodes of the default mode, dorsal attention and frontoparietal networks, onto the interaction of group x network in a linear mixed effects model, including by-subject random intercepts. We included covariates for age, sex and scan acquisition. As with the ROI analyses, we also conducted sensitivity analyses to test whether the results held when controlling for between-subjects variance in heart rate and respiration rate.

There was a significant group*network interaction for hub connectivity, such that meditators had lower participation coefficients than meditation-naïve participants in the DMN compared to other networks (*p*s<0.001 with or without physiological covariates, Figure 2a). In contrast, there was no group*network effect for within-module degree (*p*s>0.10). We further examined the effect of long-term mindfulness meditation on hub connectivity by testing group differences in participation coefficients for each network separately. Meditators’ DMN nodes had significantly lower participation coefficients compared to meditation-naïve participants’ (*F*(162)=6.02, *p*=0.02). Similar to the ROI analysis, we added heart rate and respiration rate to the model to test whether these physiological variables accounted for additional or overlapping variance in hub connectivity. While there was no significant effect of heart rate (*p*=0.51) or respiration rate (*p*=0.63) on DMN participation coefficients, above and beyond effects of the other variables, the effect of group on participation coefficients in the DMN became marginal (*F*(151)=3.62, *p*=0.06) when including these covariates. Contrary to our hypothesis, there were no differences between groups for participation coefficients in frontoparietal control or dorsal attention networks (*p*s>0.05).

**Figure 2.**
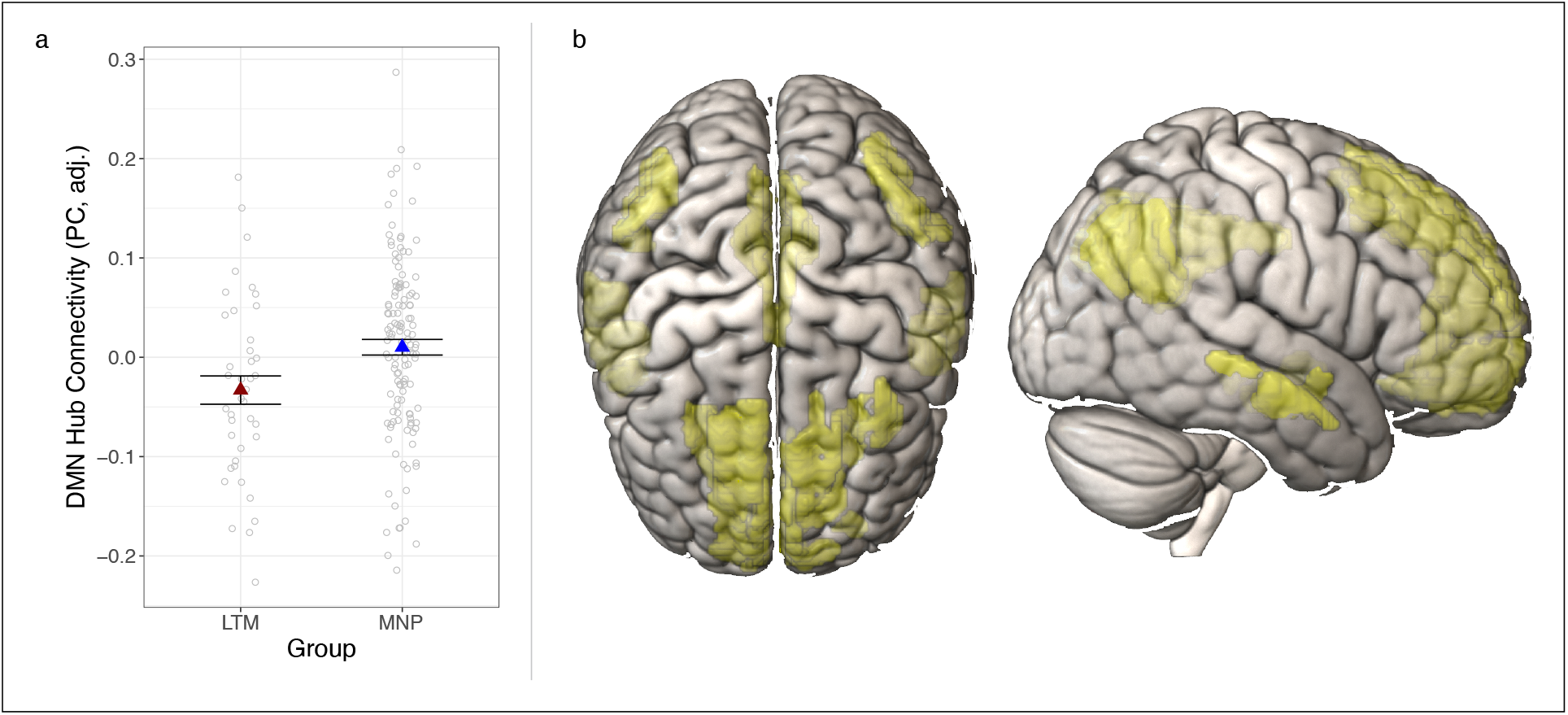
Long-term mindfulness meditation practice associated with lower DMN hub connectivity. **(a)** meditators had lower DMN hub connectivity (assessed via participation coefficients [PC]) compared to meditation-naïve participants. Dependent variables and data points are adjusted for age, gender, and scan acquisition version. Data points represent average (adjusted) PC values for each subject. Error bars represent 1 standard error above and below the point estimates of the means. **(b)** The default mode network is in yellow based on the Gordon atlas of resting state networks (Gordon et al., 2016). LTM = long term meditator; MNP = meditation-naïve participant; DMN = default mode network

### Attention

We regressed each attention measure on group (separately), while controlling for age and gender.

#### Self-report measures of attention

Meditators reported significantly greater attention to task than meditation-naïve participants, on average, during one week of experience sampling (*t*(161)=-3.42, *p*<0.01, *b*=-0.67, CI=[−1.06, −0.28]; Figure 3a). Meditators also reported higher attention than meditation-naïve participants on the ESQ (*t*(146)=-2.26, *p*=0.02, *b*=-0.40, CI=[−0.75, −0.05], Figure 3b).

**Figure 3.**
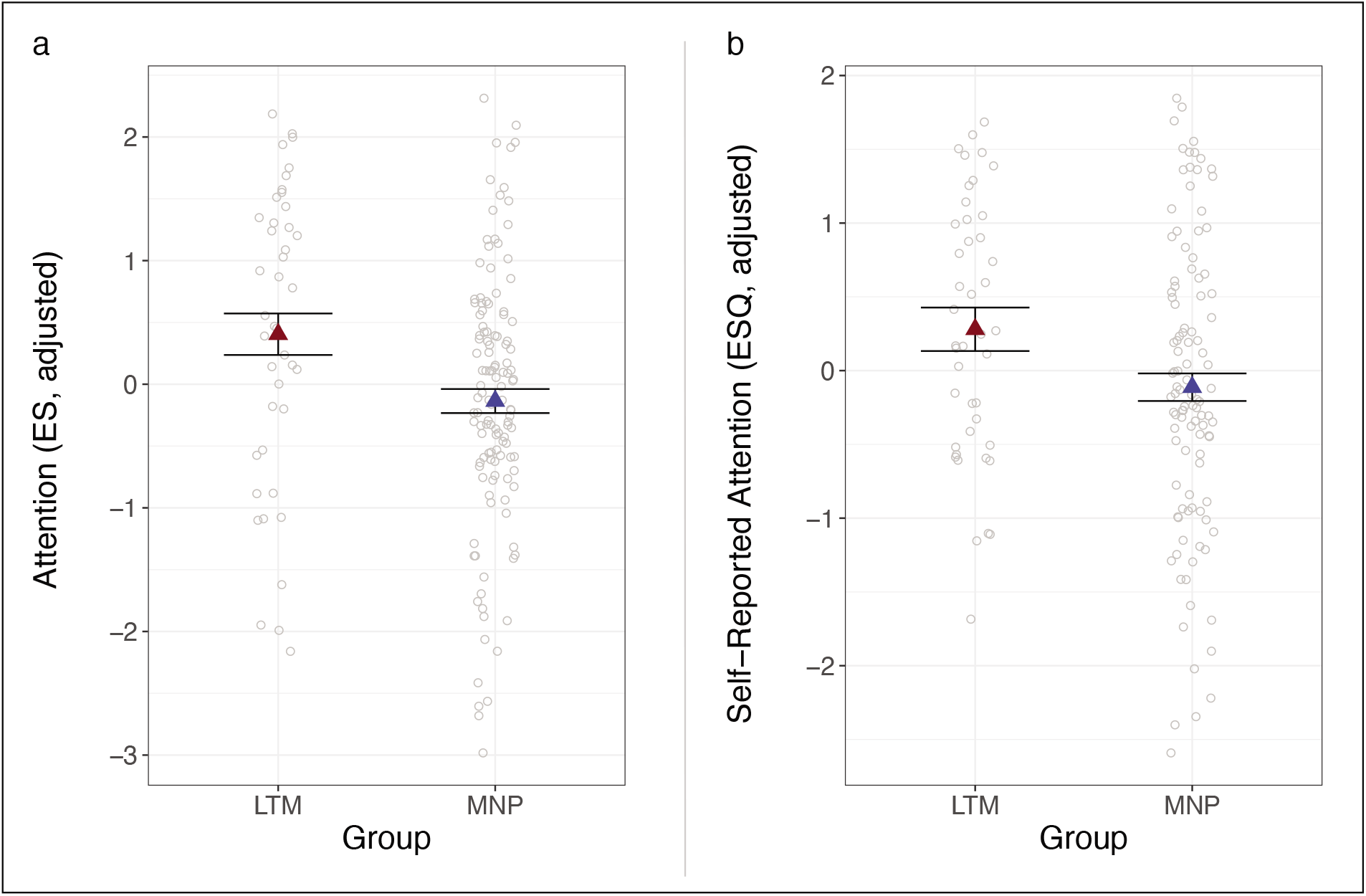
Long-term mindfulness meditation practice associated with improved self-reported attention. **(a)** Meditators had higher attention to task compared to meditation-naïve participants based on experience sampling. **(b)** Meditators had higher self-reported attention than meditation-naïve participants based on the Emotional Styles Questionnaire (ESQ) attention scale. Dependent variables and data points are adjusted for age and gender. Error bars represent 1 standard error above and below the point estimates of the means. LTM = long-term meditator; MNP = meditation-naïve participant; ES = experience sampling

#### Behavioral measures of attention: ANT

There were no differences between groups in reaction time on the ANT for orienting (t(169)=1.45, *p*=0.15, *p*\*=0.48, *b*=10.41, CI=[−3.81, 24.62]; Figure 4a), alerting (t(167)=-1.38, *p*=0.17, *p*\*=0.48, *b*=-6.43, CI=[−15.64, 2.78]), or executive control (t(165)=0.60, *p*=0.55, *p*\*=0.55, *b*=4.36, CI=[−10.01, 18.72]).

**Figure 4.**
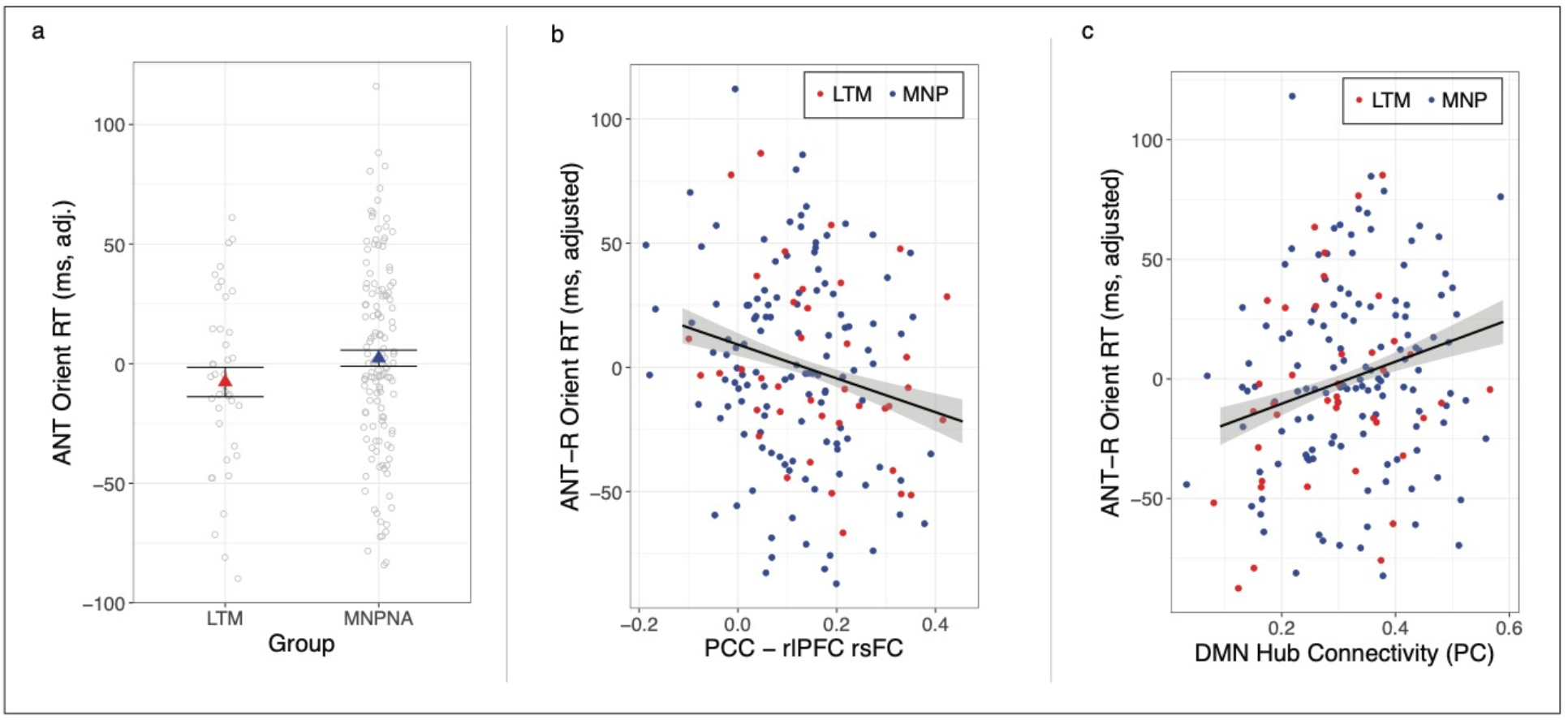
Long-term mindfulness meditation practice, orienting attention, and PCC – rlPFC rsFC. **(a)** There was no difference between meditators and meditation-naïve participants in orienting attention RT on the ANT. **(b)** Stronger PCC– rlPFC rsFC was associated with faster orienting attention, across all participants. **(c)** Lower DMN hub connectivity (assessed via participation coefficients [PC]) was associated with faster orienting attention, across participants. Dependent variables and data points are adjusted for age and gender. Error bars represent 1 standard error above and below the point estimates of the means. LTM = long-term meditator; MNP = meditation-naïve participant; PCC = posterior cingulate cortex; rlPFC = rostrolateral prefrontal cortex; rsFC = resting state functional connectivity; ANT = Attention Network Task; DMN = default mode network

### Brain – behavior relationships

In order to directly test relationships between rsFC metrics and attention, we regressed each attention measure (separately) onto rsFC, controlling for age, sex, and scan acquisition. Stronger PCC – rostrolateral PFC connectivity was associated with faster attentional orienting (t(156)=-2.77, *p*=0.01, *p*\*=0.03, *b*=-78,34, CI=[−134.13, −22.55]; Figure 4b) across all participants, and this relationship remained significant when heart and respiration rate were added to the model (*p*<0.01). Post-hoc tests showed that the relationship between stronger PCC – rostrolateral PFC connectivity and faster orienting attention was significant within the meditation-naive group (t(117)=-2.03, *p*=0.045, *b*=-72.79, CI=[−143.82, −1.75]), and marginal among meditators (t(35)=-1.78, *p*=0.08, *b*=-91.85, CI=[−196.38, 12.69]), likely due to the smaller sample size and reduced statistical power in the latter group. There was no relationship between PCC – rostrolateral PFC connectivity and alerting (t(158)=0.68, *p*=0.68, *p*\*=0.50, *b*=12.57, CI=[−23.80, 48.75]) or executive control (t(157)=1.88, *p*=0.06, *p*\*=0.09, *b*=50.87, CI=[−2.56, 104.29]).

There was no relationship between either measure of self-reported attention and PCC rsFC in the voxelwise analysis, either whole-brain or small-volume corrected to DLPFC (*p*s>0.05 corrected for FWE), nor in the ROI analysis with rostrolateral PFC (experience sampling: *t*(146)=-0.73, *p*=0.47, *p*\*=0.99, *b*=-0.61, CI=[−2.26, 1.04]; ESQ: *t*(143)=-1.05, *p*=0.30, *p*\*=0.77, *b*=-0.80, CI=[−2.32, 0.71]).

Lower average participation coefficients in the DMN were associated with faster orienting attention (*t*(164)=2.75, *p*=0.01, *p*\*=0.03, *b*=90.19, CI=[25.53, 154.86]; Figure 4c), and the result remained consistent when controlling for physiological variables (*p*<0.05). There was no relationship between DMN participation coefficients and alerting (*t*(165)=-1.73, *p*=0.09, *p*\*=0.14, *b*=-35.38, CI=[−75.80, 5.05]) or executive control (*t*(165)=-0.27, *p*=0.78, *p*\*=0.78, *b*=-9.01, CI=[−73.93, 55.91]). There was also no significant relationship between DMN participation coefficients and self-reported attention (experience sampling: *t*(152)=-0.05, *p*=0.96, *p*\*=0.96, *b*=-0.04, CI=[−1.88, 1.79]; ESQ: *t*(135)=-1.52, *p*=0.13, *p*\*=0.26, *b*=-1.38, CI=[−3.18, 0.41]).

### Practice time

There were no significant effects of practice time (either total hours of retreat or daily practice) on any brain or behavioral measures (*t*s < 2.30, *p*\*s > 0.05).

### Exploratory voxelwise connectivity analysis

In order to test for sub-regions of the DLPFC within which meditators may have had stronger PCC rsFC than meditation-naïve participants, we conducted small-volume corrected voxelwise analysis, in the bilateral, anatomically defined DLPFC mask, given the relatively large size and potential functional heterogeneity of this region. There were no statistically significant differences in PCC rsFC between meditators and meditation-naïve participants within the DLPFC mask. There were no regions in which PCC rsFC differed for meditators compared to meditation-naïve participants in wholebrain analysis, contrary to prior research (Brewer et al., 2011). Un-thresholded statistical maps are available at Neurovault (Gorgolewski et al., 2015): https://identifiers.org/neurovault.collection:8451.

## Discussion

The current study partially replicated prior work (Brewer et al., 2011), showing stronger rsFC between PCC and rostrolateral PFC in participants with a long-term practice in mindfulness meditation compared to meditation-naïve participants. Yet, we failed to replicate prior reports of stronger rsFC between PCC and DLPFC in association with long-term meditation practice. Importantly, the current study showed that stronger PCC – rostrolateral PFC rsFC was associated with faster orienting attention, indicating a functional behavioral correlate of meditation-related differences in brain connectivity of these regions. While meditators did not differ from controls in this behavioral measure of orienting attention, they reported higher goal-directed attention on both questionnaire and experience sampling measures compared to non-meditating controls. The meditation-related increases in PCC – rostrolateral PFC rsFC in the current study appear to be limited to PCC and rostrolateral PFC, as the effects were not mirrored in similar group differences in the larger brain networks to which PCC and rostrolateral PFC contribute (DMN and FPC, respectively).

Moreover, we found a relationship whereby stronger PCC – rostrolateral PFC rsFC was associated with faster attentional orienting on the ANT across groups, which lends further support for the interpretation that stronger PCC – rostrolateral PFC connectivity reflects better attention to present-moment experience. Activation of rostrolateral PFC and its connectivity with PCC could thus serve as a neural mechanism underlying subcomponents of meta-awareness that support background awareness of the source of cognition, and the ability to re-allocate attentional resources to examine internally-generated cognitions as they arise, or conversely to return attentional focus to externally-generated phenomena. While there is a wealth of research and related paradigms examining phasic instances of meta-awareness in the context of error detection and noticing instances of mind-wandering (Hester et al., 2005; Schooler et al., 2011; Ullsperger et al., 2010), research and paradigms for assessing the neural basis for sustained meta-awareness are lacking and critical for our understanding of mechanisms of change with mindfulness-based meditation training.

We also examined resting connectivity at the network level, by testing group differences in provincial (i.e., within-module) and connector (i.e., between-module) hub properties. We found *lower* participation coefficients for the default mode network in meditators compared to meditation-naïve participants, reflecting a reduction in connector hubs in the DMN. This contrasts with the stronger PCC – rostrolateral PFC rsFC we found for meditators in seed-based analysis. This may reflect reduced information flow from the default mode network to other networks, and highlights the potential specificity of increased rsFC between PCC and DLPFC in association with mindfulness meditation practice. Lower connector hub strength of the DMN was associated with *faster* orienting attention on the ANT, providing initial evidence that reduced information flow between DMN and other networks may contribute to aspects of improved attention. However, these results should be interpreted with caution, given that the effects became marginal when controlling for individual differences in heart rate and respiration rate that can confound resting state fMRI analysis. There were no differences between groups for hub properties of the dorsal attention or frontoparietal control networks, nor for provincial hub properties (i.e., within-module degree) or overall strength of connectivity between DMN and the dorsal attention network or frontoparietal control network.

Meditators had higher attention to task than meditation-naïve participants, as assessed with both a self-report questionnaire and experience sampling via text messaging. Higher attention to task, and conversely, less mind-wandering, are consistent with the goals of mindfulness meditation, namely, practicing maintaining attention on present moment experience. These behavioral results are also consistent with the interpretation of higher PCC – rostrolateral PFC resting connectivity among meditators as reflecting better attentional control of mind-wandering. However, there was no association between rsFC and self-reported attention in the current study, although we previously found that increased self-reported attention was associated with increased PCC – DLPFC rsFC following MBSR (Kral et al., 2019). It is possible that changes in these measures of attention and functional connectivity track together because they share underlying mechanisms of change, whereas the same measures examined cross-sectionally are uncorrelated due to different functionality. It is also possible that the within-subjects design in the prior RCT study may have been more sensitive to identifying correlations among changes without including variance in unrelated between-subject factors. The current study is limited given the cross-sectional nature of the data, and the associated lack of a baseline for meditators with which we could compare the current measures to examine change over time associated with long-term mindfulness meditation practice.

The rostrolateral PFC region in which we found a significant group difference was defined based on coordinates from a study showing stronger rsFC with PCC following a short-term mindfulness meditation intervention (Creswell et al., 2016). There was no difference between meditators and meditation-naïve participants in anatomically-defined ROIs, which include a more canonical DLFPC region, and in which we previously found increased rsFC with PCC following MBSR compared to controls (Kral et al., 2019). There is no overlap between the rostrolateral PFC ROI defined from the literature and the anatomically-defined DLFPC ROIs (Figure 1b, in green and light blue, respectively).

We found increased PCC rsFC in meditators relative to meditation-naïve participants in a rostrolateral PFC ROI that is situated in Brodmann’s Area 10. While this region is activated across numerous cognitive tasks, taken together, the available evidence lends support to the hypothesis that rostrolateral PFC may switch attention between internal and external stimlui (e.g., between self-related processing and task- or goal-oriented processing (Burgess et al., 2007). According to this hypothesis, rostrolateral PFC is part of a system that serves to determine the source of cognitive representations (e.g., internally or externally generated), exerting influence on attention allocation in open-ended situations when goals are self-generated or underspecified (e.g., the task-free setting of a resting state scan), or when sustained attention is required (Burgess et al., 2007). This hypothesized function for rostrolateral PFC is consistent with a role in meta-cognitive background awareness that is a core component of mindfulness meditation training. This form of background meta-awareness is specifically cultivated in “open monitoring” style practices that emphasize broad awareness to present-moment experience while simultaneously monitoring for the presence of mind-wandering, and returning attention to the “task” of focusing on present-moment experience (Lutz et al., 2015).

The results of the current study provide additional evidence consistent with prior work indicating a relationship between mindfulness meditation practice and increased resting connectivity between nodes of the default mode and frontoparietal control networks (Brewer et al., 2011; Creswell et al., 2016; Kral et al., 2019). We also found evidence for a relationship between default mode network – frontoparietal network functional connectivity at rest and attentional orienting, and importantly, the meditators had higher goal-directed attention than meditation-naïve participants in experience sampling and self-report measures. Changes in the dynamic interactions of PCC and rostrolateral PFC are a candidate neural mechanism underlying attentional improvements with mindfulness meditation training. Further research is needed to determine the mechanisms and trajectory of change with mindfulness meditation practice, including contextual and individual differences factors that may influence its efficacy.

## Methods

This study is registered as a clinical trial with ClinicalTrials.gov (NCT02157766). Some of the methods detailed below were previously described (Kral et al., 2019).

### Participants

We recruited 183 healthy participants from a non-clinical population, comprised of 140 meditation-naïve participants and 43 meditators. Meditation-naïve participants (average age 44.3±12.8 years, 83 female) were recruited from Madison, WI and the surrounding community using flyers, online advertisements, and advertisements in local media for a study researching “the impact of health wellness classes on the brain and body”. The baseline data for meditation-naïve participants (participants in the RCT) served as a control group in this cross-sectional study. Seventeen meditation-naïve participants had rsFC data excluded from analysis due to excessive motion (described below; n=11) or anatomical brain abnormalities as determined by a neuroradiologist (n=6), resulting in inclusion of 123 meditation-naïve participants (average age ± SD = 42.4 ± 12.4 years, 74 female, 49 male) in analyses reported here.

Meditators were recruited from meditation centers and through related mailing lists throughout the United States, in addition to flyers and advertisements in newspapers similar to the recruitment strategy for meditation-naïve participants. Meditation-related recruitment criteria included at least five years of daily practice (with an average practice of at least 200 minutes per week), experience with Vipassana, concentration and compassion/loving-kindness meditations, and at least 5 weeks of retreat practice. Meditation retreats involve spending a continuous period of days or weeks (and in some cases, years) in meditation practice, often at a meditation or community center. Retreats often include group practice, in addition to solitary practice, and may include extended periods of silence. Lifetime hours of practice were calculated based on participants’ reports of their average hours of formal meditation practice per week and their total years of practice (average = 3759 hours, range = 780 to 19,656 hours). Lifetime retreat practice hours were calculated by summing the practice hours that were reported for each retreat. Practice hours were log-transformed using the natural log, to correct for a highly right-skewed distribution. Three meditators had rsFC data excluded from analysis due to excessive motion (n=2) or anatomical brain abnormalities (n=1), resulting in 40 meditators in the final sample for analysis (average age ± SD = 44.1±11.8 years, 15 female, 25 male).

Participants were excluded if any of the following applied, due to their potential impact on the current analyses or other aspects of the larger study in which they were enrolled: regular use of psychotropic or nervous system altering medication; psychiatric diagnosis in the past year or history of bipolar disorder, schizophrenia or schizoaffective disorder; color blindness; currently participating in another clinical trial (meditation-naïve participants only); current asthma diagnosis; currently diagnosed with a sleep disorder or regularly taking prescribed sleeping medications; current night shift worker; significant training or practice in meditation or mind-body techniques such as yoga or Tai-Chi (meditation-naïve participants only); expert in physical activity, music or nutrition (meditation-naïve participants only); any history of brain damage or seizures; medical conditions that would affect the participant’s ability to participate in study procedures. Written, informed consent was obtained from all participants according to the Declaration of Helsinki (“WMA - The World Medical Association-WMA Declaration of Helsinki – Ethical Principles for Medical Research Involving Human Subjects,” n.d.) and the study was approved by the Health Sciences Institutional Review Board at the University of Wisconsin–Madison.

### Data collection

Participants attended a 24-hour lab visit at the Waisman Laboratory for Brain Imaging and Behavior that included an MRI scan, behavioral testing, self-report data collection, and additional measures as part of a larger multi-session, multi-project study. Experimenters were blind to the group assignment during data collection. All participants were given monetary compensation for their participation.

#### Emotional Style Questionnaire

The Emotional Style Questionnaire (ESQ; Kesiber et al., 2019) consists of a 1 – 7 Likert scale with 1 = strongly disagree and 7 = strongly agree. One of the ESQ sub-scales provided a measure of attention that was most relevant to the hypotheses of the current study, and items included: “I do not get distracted easily, even when I am in a situation in which a lot is going on” and “I sometimes feel like I have very little control over where my attention goes” (reverse-coded). A subset of meditation-naïve participants(n=93, average age ± SD = 44.1±11.8 years, 48 female, 45 male) completed the ESQ, which was introduced subsequent to the onset of data collection due to availability of the measure. All meditators completed the ESQ.

#### Attention Network Task (ANT)

The ANT was performed outside the scanner. Stimuli were presented using E-prime 2.0 (*E-Prime 2.0*, 2012), and the task was administered as described in prior literature (Fan et al., 2009). We calculated reaction time (RT) for each condition after excluding error trials, and RTs from trials longer than 1200 ms (errors of omission) and shorter than 250 ms (impulsive responses) were also excluded. Two participants were excluded due to low accuracy (one participant had 18% accuracy on congruent trials; and one participant had 51% accuracy on incongruent trials).

#### MRI acquisition

Images were acquired on a GE MR750 3.0 Tesla MRI scanner with a 32-channel head coil. Anatomical scans consisted of a high-resolution 3D T1-weighted inversion recovery fast gradient echo image (inversion time = 450 msec, 256×256 in-plane resolution, 256 mm FOV, 192×1.0 mm axial slices). A 12-minute functional resting state scan run was acquired using a gradient echo EPI sequence (360 volumes, TR/TE/Flip = 2000 ms/20 ms/75°, 224 mm FOV, 64×64 matrix, 3.5×3.5 mm in-plane resolution, 44 interleaved sagittal slices, 3-mm slice thickness with 0.5 mm gap). The in-plane resolution was decreased after the first 21 participants from 3.5×3.5 mm to 2.33*3.5 mm to better address sinus-related artifacts, resulting in a matrix of 96×64.

#### Experience sampling

Experience Sampling was conducted for one week following the lab visit. Participants provided their mobile phone numbers and availability for 8-hour periods for each of the 7 days. Participants had a choice of receiving text messages 6, 7, or 8 times a day, and received a text message every 90 minutes on average. The text message contained a question assessing mind-wandering: “Was your attention on the activity you were performing?” Participants were asked to respond with a number from 1 (attention is not on the task) to 9 (attention is completely on the task at hand). On average participants responded to 82% of text messages they received. The response window was set to the time between two successive messages, such that participants were given until the next message arrived to respond to the current message. If participants sent two responses in-between messages, the second response was discarded. The ratings across all 7 days of the week were averaged to obtain a mean attention rating for each participant.

### Data Analysis

#### Functional image processing & analysis

Functional images were processed using a combination of AFNI (Cox, 1996) version 17.3 and FEAT (FMRI Expert Analysis Tool) Version 6.00, part of FSL (FMRIB’s Software Library) (Smith et al., 2004), using the following steps: removal of the first 4 volumes; motion correction with MCFLIRT (Jenkinson et al., 2002); brain extraction with BET (Smith, 2002); registration of the subject’s functional data to their anatomical image using the Boundary-Based Registration approach (Greve and Fischl, 2009). A 12-degree of freedom affine transformation using FLIRT (Jenkinson et al., 2002) was followed by FNIRT nonlinear transformation to register each subject’s functional data to Montreal Neurological Institute 152 space. Images were segmented into white matter, grey matter and cerebrospinal fluid (CSF) with FAST for use as masks that were eroded using a 3×3×3 voxel kernel and then used to generate ROI-averaged time series, with white matter and CSF time-series serving as nuisance regressors (along with their derivatives and the 6 motion regressors) with AFNI’s 3dDeconvolve. Images were smoothed with a 5-mm full-width half-maximum Gaussian kernel.

We extracted the time-series from a spherical PCC seed with a 4-mm radius defined based on coordinates from Creswell *et al*. (2016) (Figure 1b, in yellow). We regressed each time-series (separately) back onto each subject’s data using AFNI’s 3dDeconvolve, which also censored high-motion time-points (greater than 0.2 mm framewise displacement) (Power et al., 2014). Participants were excluded from analysis if they had less than 6 minutes of data due to more than 50% of data points censored for motion. Two sets of target ROIs were defined for assessing PCC rsFC: a bilateral DLPFC ROI, based on medial frontal gyrus (MFG) from the Harvard-Oxford atlas (Craddock et al., 2012) thresholded at 50% probability for small-volume-corrected voxelwise analysis (Figure 1b, in light blue), which was split into left and right for ROI analysis; and a left rostrolateral PFC ROI defined as a 10-mm sphere around coordinates provided in Creswell *et al*. (2016) (Figure 1b, in green). Connectivity was assessed based on the Fisher-Z transformed correlation between the seed and every other voxel in the brain for the voxelwise analysis, and separately for each of the target ROIs. Voxelwise analyses were thresholded at *p*<0.05 controlling for family-wise error using threshold-free cluster enhancement with FSL’s Randomise (Winkler et al., 2014).

#### Graph theoretical network analysis

We calculated hub connectivity metrics for the default mode (Figure 2b), frontoparietal control, and dorsal attention networks based on the Gordon connectivity atlas (Gordon et al., 2016). First, the mean resting state time-series was extracted from each of the 333 nodes in the Gordon atlas, and then we constructed a correlation matrix for each subject by computing pairwise Pearson correlations for each set of nodes. We used the correlation matrix to calculate both provincial and connector hub properties of each node using within-module degree (WMD) and participation coefficient measures, respectively, (with nodes assigned to networks as defined in the Gordon atlas) following the procedures detailed by Hwang, Bertolero, Liu & D’Esposito (Hwang et al., 2017). We tested for group differences in hub connectivity across all nodes in the respective network (default mode, frontoparietal control, dorsal attention or salience network), separately, by estimating linear mixed-effects models with the lmer and anova functions in R statistics (Kuznetsova et al., 2017; R Core Team, 2013), which included by-subject random effects and covariates to control for age, gender and scan acquisition version.

#### Physiological measures

Respiration and heart rate data were collected during the concurrent resting state fMRI scan, amplified using a BIOPAC MP-150 system, and digitized at 1000 Hz. Mean heart rate was assessed using CMetX (Hibbert et al., 2012) and inter-beat interval (IBI) series were corrected for artifact and ectopic beats. Heart rate data were cleaned by interpolating over ectopic or artifactual IBI using inhouse MATLAB software (Allen et al., 2007). Respiration data were collected using a respiration belt placed over the thorax, data were inspected for artifact, and artifacts were rejected. Mean respiration rate was calculated with in-house Matlab scripts using trough-to-trough measurements on the cleaned respiration series.

#### Statistical analysis

Linear models were conducted using the lm function in the stats package in R (R Core Team, 2015), in which rsFC (or other dependent variable) was regressed on group (or other variables of interest). All analyses included covariates for age and sex, and analyses of rsFC included an additional covariate for the change in the resting state scan acquisition (as described above). All results are reported after removing outliers based on Cook’s D, with a cutoff threshold of 4/(N-P) for data points disconnected from the distribution (where N=sample size and P=number of parameters in the model) as determined by the modelCaseAnalysis function of the lmSupport package (Curtin, 2015) in R (R Core Team, 2015). The number of outliers removed from each analysis ranged from 0 to 5 meditation-naïve participants, and from 0 to 2 meditators. We used a false discovery rate (FDR) correction to control for multiple comparisons for each family of tests (e.g., across 3 ROIs), using the p.adjust function in R statistics, and corrected *p*-values are indicated by *p*\*.

## Acknowledgments

This work was supported by the National Center for Complementary and Integrative Health (NCCIH) [P01AT004952] to RJD, grants from the National Institute of Mental Health (NIMH) [R01-MH43454, P50-MH084051] to RJD, grants from the Fetzer Institute [2407] and the John Templeton Foundation [21337] to RJD, a core grant to the Waisman Center from the National Institute of Child Health and Human Development [P30 HD003352-449015] to Albee Messing. TRAK was supported by the National Institute of Mental Health award number T32MH018931.

We would like to thank Michael Anderle, Ron Fisher, Jane Sachs, Jeanne Harris, Mariah Brown, Elizabeth Nord, Kaley Ellis, Gina Bednarek, Kara Chung, Pema Lhamo, David Bachhuber, Amelia Cayo, Christopher Harty, Sonam Kindy and Dan Dewitz for assistance with recruitment and/or data collection. We would also like to thank Katherine Bonus, Devin Coogan, Bob Gillespie, Diana Grove, Lori Gustafson, Matthew Hirshberg, Peggy Kalscheur, Chad McGehee, Vincent Minichiello, Laura Pinger, Lisa Thomas Prince, Kristi Rietz, Sara Shatz, Chris Smith, Heather Sorensen, Jude Sullivan, Julie Thurlow, Michael Waupoose, Sandy Wojtal-Weber and Pam Young for teaching and/or coordinating the interventions, and John Koger, Ty Christian, David Thompson and Nate Vack for technical assistance.

## Author Contributions

T.I. contributed to data collection. T.R.A.K., T.I., E.P., and D.W.G. processed the data. T.R.A.K. analyzed the data and wrote the manuscript. T.I., R.L., and E.P. contributed to planning the analysis. R.L., R.G., and M.A.R. contributed to the design of the study. R.J.D. designed and supervised the study. All authors contributed to writing the manuscript and approved it for submission.

